# Phylogeography by Diffusion on a Sphere

**DOI:** 10.1101/016311

**Authors:** Remco R. Bouckaert

**Affiliations:** Computational Evolution Group, University of Auckland Computer Science Department, University of Waikato Institute for History and the Sciences, Max Planck Institute

## Abstract

Techniques for reconstructing geographical history along a phylogeny can answer many interesting question about the geographical origins of species. Bayesian models based on the assumption that taxa move through a diffusion process have found many applications. However, these methods rely on diffusion processes on a plane, and do not take the spherical nature of our planet in account. This makes it hard to perform analysis that cover the whole world, or do not take in account the distortions caused by projections like the Mercator projection.

In this paper, we introduce a Bayesian phylogeographical method based on diffusion on a sphere. When the area where taxa are sampled from is small, a sphere can be approximated by a plane and the model results in the same inferences as with models using diffusion on a plane. For taxa samples from the whole world, we obtain substantial differences. We present an efficient algorithm for performing inference in an Markov Chain Monte Carlo (MCMC) algorithm, and show applications to small and large samples areas.

Availability: The method is implemented in the GEO_SPHERE package in BEAST 2, which is open source licensed under LGPL.

## 1 Introduction

A number of Bayesian phylogeographical methods have been developed in the recent years [16, 17, 6, 20] that make it convenient to analyse sequence data associated with leaf nodes on a tree and infer the geographical origins of these taxa. This allows for answering questions about geographical origins of for example viral outbreaks like Ebola [12], and HIV [11], the Indo-European language family [6] as well as many other species (see [10] for more).

The assumption underlying these methods is that taxa migrate through a random walk over a plane. This may be appropriate for smaller areas, but when samples are taken from a large area of the planet, it may be necessary to map part of the planet onto a plane. Figure 1 shows two such projections: the popular Mercator projection, which shows large distortions in areas especially around the poles and the Mollweide projection which ensures equal areas, but distorts the relative positions. Though some projections preserve some metric properties, unfortunately, there is no projection that maintains distances between all pairs of points.

**Figure 1:**
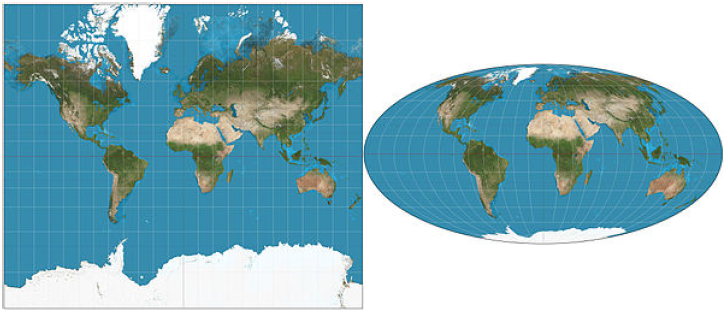
Left Mercator projection, right Mollweide projection.

In this paper, we consider random walks over a sphere, instead of over a plane. This is equivalent to assuming a heterogeneous diffusion process over a sphere. The benefit is that we do not need to worry about distortions in the projection. Also, for smaller areas it behaves equivalent to a model assuming diffusion over a plane, so it can be applied on any scale. Perhaps the most closely related model is described in [9], which uses a less accurate approximation for spherical diffusion and does not work out an efficient inference scheme.

In the next section, we detail the model, which follows the tradition of treating the geography as just another piece of information on each of the leaves in the tree. In this model, the geography is independent of any sequence information for the leaves conditioned on the tree. Section 3 has implementation details for using the model efficiently with MCMC. We develop an approximate likelihood that can be calculated efficiently and relies on setting the internal locations to their (weighted) mean values. We compare results between planar and spherical diffusion in a simple simulation study and analyse the origin of Hepatitis B in Section 4, then wrap up with a discussion and conclusions.

## 2 The spherical diffusion model

In this section, we explain the details of the spherical diffusion model and how to apply it to phylogeography. We assume homogeneous diffusion over a sphere, governed by a single parameter, the precision *D* of the diffusion process. For such diffusion, the probability density of making a move over angle at time *t* is closely approximated by [18]:

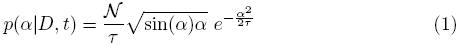

where *𝒩* is a normalising condition such that 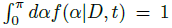 and τ = 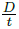. Figure 2 shows the shape of *𝒩* for different values of *τ* calculated using numeric integration in R. For small values of *τ*(< 10^−6^) we found that *𝒩* is approximately 1 (> 0.9999999), while for large values of *τ*(> 10^6^) *𝒩* approaches 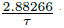 within a relative error bounded by 10^−6^. In between we can pre-calculate the values of *τ* in the range 2^*a* /2^ for *a* ∈ {−40, 60} and get reasonably close values by interpolating between these samples, since the function of *𝒩* in *τ* is very smooth.

**Figure 2:**
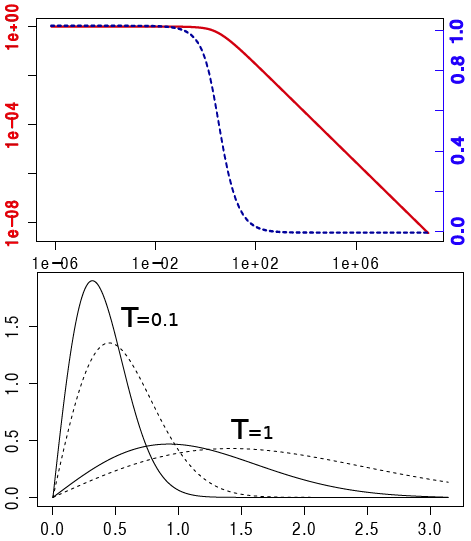
Topt, 1/*𝒩*(*τ*) (y-axis) for different values of *τ* = *D/t* (x-axis) on a log-log scale (left axis, solid line) and normal-log scale (right axis, dashed line). Bottom, density functions (y-axis) for spherical diffusion (solid lines) and planar diffusion (dashed lines) with *τ* = 0.1 and *τ* = 1.0. The x-axis is the angle for spherical diffusion and distance to start point for planar diffusion.

Figure 2 shows the difference between a density function with precision 1 and at time 1 for both planar diffusion and spherical diffusion. Spherical diffusion peaks a bit earlier, but planar diffusion has a longer tail. This intuitively makes sense; there is less space on a sphere when moving out 1 unit than on a plane, so a slightly earlier peak is expected. Also, for a long distances on a sphere on arrives on the other side at a smaller angel, once the sphere is traversed to the other side.

Let *T* be a binary tree over a set of *n* taxa *x* _1_,…,*x*_*n*_. Internal nodes of the tree are numbered *n* + 1,…, 2*n −* 1. With each node *x*_*i*_ is associated a location (*lat*_*i*_, *long*_*i*_) with latitude *lat*_*i*_ and longitude *long*_*i*_.

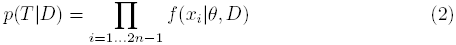

where *D* the precision of the diffusion process and *θ* other parameters, like branch length, etc., and *f* (*x*_*i*_|,*θ*,*D*) defined as

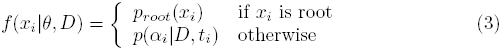

with *p*_*root*_ (.) the root density, and *p* (*α*_*i*_|*D*, *t*_*i*_) the spherical density of Equation 1 and *α*_*i*_ the angle between location of *x*_*i*_ and its parent, and *t*_*i*_ the ’length’ of the branch. This length is equal to the clock rate used for the branch times the length of the branch in the tree. For instance for a strict clock with rate *r*, the length is simply equal to the branch length in time times *r*. More relaxed models like the uncorrelated relaxed clock [8], have individual rates that can differ for each branch.

For the root density *p*_*root*_ (.), we usually take the uniform prior, indicating we have no preference where the root locations is placed. This simplifies Equation (2) in that the first term becomes a constant, which can be ignored during MCMC sampling. Using more complex priors can have consequence for inference, as outlined in Section 3.3.

To determine the angle between two locations (*lat*_1_*, long*_1_) and (*lat*_2_*, long*_2_)requires a bit of based geometry:

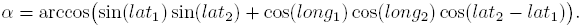

A common situation is where we have a tree where the leaf nodes have known point loctions and we want to infer the locations of internal nodes, in particular the root location which represents the origin of all taxa associated with leaf nodes in the tree. The tree is typically informed by sequence information *D* associated with the leaf nodes.

## 3 Implementation

The simplest, and in many ways most flexible, approach is to explicitly maintain all locations as part of the state. Calculation of the likelihood for geography on a tree using Equation 2 (up to a constant) becomes straightforward. Unfortunately, designing MCMC proposals that efficiently sample the state space is hard.

### 3.1 Particle filter approach

The particle filter method [7] is another way of calculating the likelihood without explicit representation of the locations. For every sample in the MCMC chain, the likelihood is calculated based on the tree as informed by other data. As a result, the MCMC has a lot lower change of getting stuck in local maxima and less care is required in designing efficient proposals for dealing with the geography.

To calculate the likelihood, we use a number of ‘particles’ and each particle represents the locations of each of the nodes in the tree. Locations of internal nodes are initialised by setting them to the mean locations of their children in a post-order traversal of the tree, which results in a sufficiently good fit of locations to the tree to guarantee quick convergence. Next, particles are perturbed in pre-order traversal as follows: for a location of node *x*_*i*_, randomly *k* locations are sampled in the vicinity of the current location. Out of the *k* locations, one location is sampled proportional to the partial fit of the location. This partial fit is simply the contribution the location provides to the density (Equation (2)) consisting of

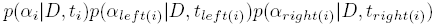

where *x*_*left* (*i*)_ is the left child of *x*_*i*_ and *xright* (*i*) its right child. For the root node *p*(α_*i*_|*D, t*_*i*_) is assumed to be constant.

After all particles are perturbed, an equally sized set of particles is sampled (with replacement) from the current set, with probability proportional to the density as defined by Equation (2). This process quickly converges to a stable likelihood. Furthermore, it does not easily get stuck in local maxima. Unfortunately, in the context of the MCMC algorithm where the likelihood needs to be recalculated many times, this process is still rather slow.

Therefore, we explore an approximation based on assigning mean locations to each of the internal nodes.

### 3.2 Mean location approximation

Suppose we want to calculate the mean location for each internal node defined as

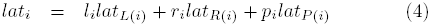

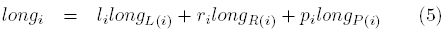

and at the root node as

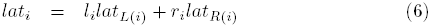

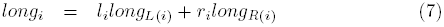

where *L*(*i*), *R*(*i*) and *P*(*i*) the index of left and right child and parent of *x*_*i*_ respectively and *l*_*i*_, *r*_*i*_ and *p*_*i*_ are positive weights associated with first and second child and parent respectively that add to unity (*l*_*i*_ + *r*_*i*_ + *p*_*i*_ = 1).

**Figure 3:**
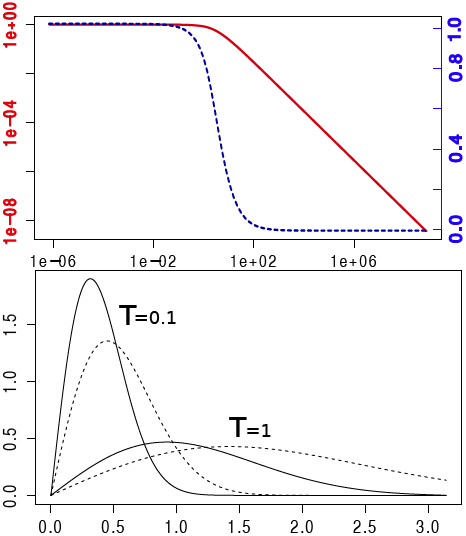
Left, the mean of two blue points by taking the mean latitude/longitude is the red point, while the green point is closer. Right, taking the mean location of 2 and 3 points on a sphere.

So, we have a set of *n −* 1 linear equations in *n −* 1 unknown values for latitude, and the same for longitude. We can solve these with standard methods for solving linear equations such as Gaussian elimination in *O*(*n* ^3^), but the structure of the tree allows us to solve this problem more in linear time (*O*(*n*)) as follows.

First, do a post order traversal where for each node, we send a message (*m*_*i*_, *ρ*_*i*_) to the parent where if *x*_*i*_ is a leaf node,

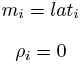

and if *x*_*i*_ is not a leaf node,

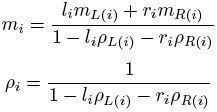

At the root, we have

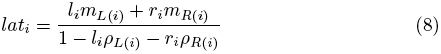

Next, we do a pre-order traversal from the root, sending down the latitude, and calculate for all internal nodes (but not leaf nodes)

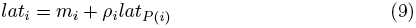

#### Theorem 1.

*Using the above calculations lat*_*i*_ *is as defined in Equations (4) and (6) and is calculated in O* (*n*).

**Figure 4:**
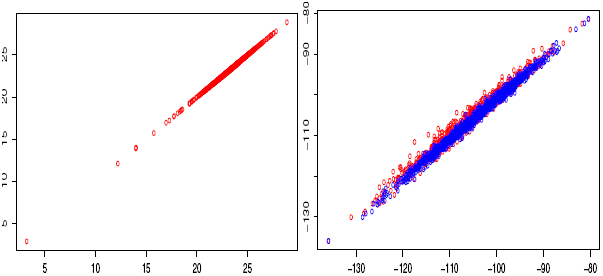
Particle filter (horizontal) vs mean approximation (vertical) on a log of log-likelihood scale. Left, sampled from prior, right sampled from posterior (in blue). Also right particle filter vs particle filter (in red) to show stochasticity of particle filter.

See Appendix A for a proof.

Especially close to the poles, taking the average point between to pairs of latitude/longitude pairs can be a point that is far from the point with the shortest distance to these points. For example the points (45, 0) and (45, 180)(blue points in Figure 3) on opposite sides of the pole have a mean of (45, 90) (red point)– a point at the same latitude – while the pole (90, 0) (green point) has a shorter distance to these two points.

Instead of using latitude/longitude to represent location, we can use Cartesian coordinates (*x, y, z*) on a sphere. We can convert (*lat, long*) locations to Cartesian using (*x, y, z*) = (sin(*θ*) cos(*ϕ*), sin(*θ*) sin(*ϕ*), cos(*θ*)) where *θ* =*longπ/* 180 and *ϕ* = (90 *− lat*) *π/* 180. To convert a Cartesian point back to latitude/longitude, we use (*lat, long*) = (arccos(*−z*)180*/π−* 90, arctan 2(*y, x*)180*/π*).

However, the mean of two points on a sphere is not necessarily on a sphere. However, if we take the mean of two points and project it onto a sphere by taking the intersection of the sphere with a line through the origin and the mean point, we get the point that has shortest distance to both points.

Figure 4 shows the likelihood calculated through the particle filter approximation and the mean location approximation outlined above. When sampling from the prior, the correlation is almost perfect, while when sampling from the posterior, the mean approximation shows a slight bias. However, also shown in Figure 4 is how well the particle filter correlates with itself. Due to the stochastic nature of the algorithm, this correlation is not 100%, and only slightly better than the correlation with the mean approximation approach.

In summary, we can use the mean approximation to determine locations of internal nodes using a fast *O*(*n*) algorithm and use Equation 2 with these locations to approximate the likelihood. The only geography specific parameter to be sampled by the MCMC is the precision parameter for the diffusion process. To log locations so that we can reconstruct the geography, logging the mean approximations would result in biased estimates of the uncertainty in locations. To prevent this, the particle filter is run for those samples that are logged and one of the particles containing all internal locations is used as representative sample for a particular state of the MCMC. Since the particle filter only needs to be run when a state is logged, this is sufficiently computationally efficient to be practical.

### 3.3 Refinements

The mean approximation outlined in the previous subsection assumes tips have fixed positions and all internal nodes have not. If tips are not point locations, but sampled from a region with known boundaries, for example a country or province, the tips can be sampled using a uniform prior over the region it is known to be sampled from. A random walk proposal for tips can be used to sample tip locations. The mean approximation runs as before, but obviously when a tip is updated, the tip location needs to be fixed at the new location for the algorithm.

Suppose a uniform prior for the root is not appropriate, but a region is known to which the root can be confined. The mean-approximation will assign a value to the root location without concern for such prior and it may assign a location to the root outside the known region. However, if the root location is represented explicitly as part of the state for the MCMC algorithm, the mean approximation can use that location in its likelihood calculation instead the one represented by Equations (6) and (7). Like tip locations, the root location can be sampled. A distribution representing whether the root locations is in the region can be added to the prior.

The same technique can be applied if the region of a clade represented by a set of taxa is known. Such location can be explicitly modeled in the state and the mean approximation, instead of using Equations (4) and (5).

## 4 Results

To compare the planar and spherical diffusion models, we run an MCMC analysis with both models using a fixed tree with three taxa that are positioned on a sphere around the location (0, 0) (see Figure 5). The same coordinates were then placed on the sphere, the sphere rotated towards the south pole in steps of 10 degrees up to 80 degrees and resulting latitude and longitude positions used for A, B and C. One would expect when inferring the root locations (R in Figure 5) that rotating it back with the same angle would result in inferring the same root location as for the unrotated problem. So, the great circle distance between the taxa A, B and C does not change when rotated.

Table 1 shows the difference in original root and unrotated root. The planar diffusion shows a significant difference in the estimate of the latitude, while it does not for the spherical diffusion estimate. The spherical diffusion model shows smaller standard deviation, possibly due to a different parameterisation than the planar model. Though both models contain the original root location in the 95% highest probability density (HPD) interval, the spherical diffusion model has a considerable lower bias than the planar model.

**Figure 5:**
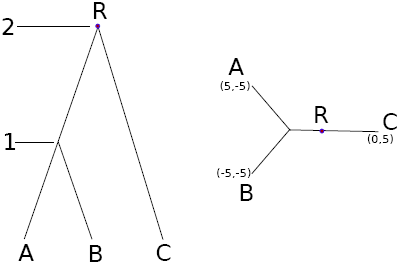
Tree ((A:1,B:1):1,C:2) and locations in latitude, longitude pairs for simulation study.

**Table 1:** Difference in root location between unrotated and rotated cases for planar and spherical diffusion models. The number after the ± are the standard error in the estimate of the mean of the number before the ±.

| Angle | Diffusion on plane |  |  |  | Diffusion on sphere |  |  |  |
| --- | --- | --- | --- | --- | --- | --- | --- | --- |
| | $\Delta$ latitude | | $\Delta$ longitude | | $\Delta$ latitude | | $\Delta$ longitude | |
| 0 | 0 | $\pm 0.19$ | 0 | $\pm 0.13$ | 0.00 | $\pm 0.06$ | 0.00 | $\pm 0.04$ |
| 10.00 | 0.16 | $\pm 0.20$ | -0.37 | $\pm 0.21$ | 0.04 | $\pm 0.05$ | -0.02 | $\pm 0.04$ |
| 20 | 0.39 | $\pm 0.25$ | -0.27 | $\pm 0.19$ | -0.11 | $\pm 0.05$ | 0.03 | $\pm 0.04$ |
| 30 | 0.66 | $\pm 0.21$ | -0.07 | $\pm 0.16$ | 0.04 | $\pm 0.05$ | 0.08 | $\pm 0.04$ |
| 40 | 1.56 | $\pm 0.30$ | -0.28 | $\pm 0.20$ | -0.04 | $\pm 0.05$ | 0.11 | $\pm 0.04$ |
| 50 | 2.36 | $\pm 0.43$ | -0.47 | $\pm 0.32$ | 0.04 | $\pm 0.06$ | 0.14 | $\pm 0.05$ |
| 60 | 3.06 | $\pm 0.38$ | -0.59 | $\pm 0.63$ | 0.03 | $\pm 0.05$ | 0.28 | $\pm 0.05$ |
| 70 | 3.96 | $\pm 0.50$ | -0.45 | $\pm 0.88$ | 0.11 | $\pm 0.06$ | 0.47 | $\pm 0.06$ |
| 80 | 4.32 | $\pm 0.39$ | 0.49 | $\pm 1.61$ | 0.18 | $\pm 0.05$ | 1.02 | $\pm 0.08$ |

**Figure 6:**
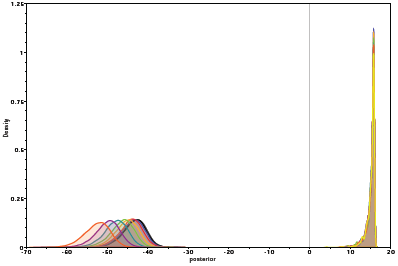
Posterior densities for planar (left side) and spherical (right side) for simulation.

The planar diffusion model has different likelihoods and posteriors, while the spherical diffusion model’s posterior is invariant under rotation of the taxa. This is illustrated by Figure 6 that shows all posteriors. The ones for planar diffusion are clearly distinguishable while the ones for spherical diffusion are too similar to be separated.

To get an impression of the capability of the spherical model, HBV full genome sequences were taken from Genbank (see Appendix B for accession numbers, sample dates used and country of origin) and clustalx [15] was used to align the sequences. Samples come from Cambodia, China, DR Congo, France, Germany, Ghana, Greece, India, Indonesia, Italy, Japan, Korea, Myanmar, Namibia, Philippines, Russia, Thailand, Turkey and Vietnam. An initial run tips were not sampled but just taken from a centroid of the country. In this case, the Russian samples clearly reside in different parts of the tree and as a consequence when visualising the tree through space, branches from both Europe and East Asia converged into the point representing Russian samples. Since such a scenario is unlikely, and it is not possible to get more accurate sample location information, tips were sampled from the regions defined by the country of origin as listed in Appendix B. Border data was obtained from http://thematicmapping.org/downloads/world_borders.php which is available under the Creative Commons license and converted to KML files at http://www.mapsdata.co.uk/online-file-converter/.

Many attempts at estimating the rate for HBV have been made (see, e.g. [4, 19, 13]) but most are inconsistent. In order to concentrate on geography and hopefully without stirring any more controversy, the clock rate was fixed at 2.0E-5 substitutions/site/year, but results can be scaled to what the reader deems more appropriate.

**Figure 7:**
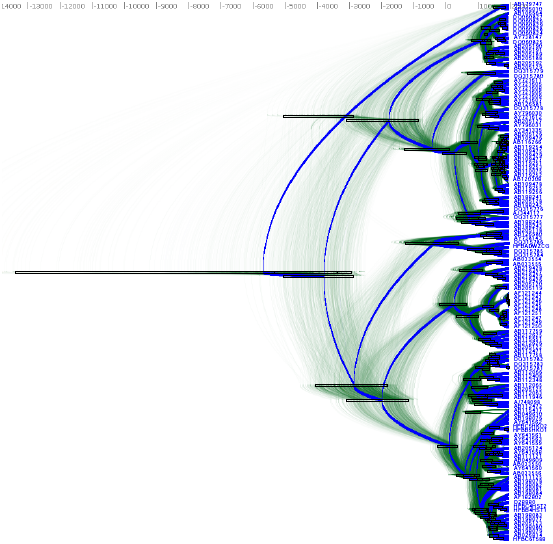
DensiTree of Hepatitis B in Eurasia and Africa.

We used BEAST 2 [5] to perform the analysis with the GTR substitution model with gamma rate heterogeneity with 4 categories and invariant sites. The uncorrelated relaxed clock with log normal distribution [8] was used and shows a coefficient of variation with of mean 0.476 and 95% HPD Interval of (0.3721, 0.5715), which means a strict clock can be ruled out. A coalescent with constant population size was used as tree prior. The spherical model was run with a strict clock and a relaxed log normal clock. The AICM values [1] of the strict clock was 69215 *±* 0.93 while that of the relaxed clock was 69086 *±* 2.063 thus favouring the relaxed clock with a difference of over 128.

Figure 7 shows the DensiTree [3] of the HBV analysis, which demonstrates it is fairly well resolved, with many clades having 100% posterior support.

Figure 8 shows the tree mapped onto the earth after processing with SPREAD [2] and visualised with google-earth. The root of the tree and thus the associated origin of the virus is placed in northern India about 10000 year ago. Figure 8 shows the spread at times -4000 CE, -1000 CE, 1000 CE and present. Also shown is a heat map visualising the internal node positions of the trees in tree set representing the posterior. Colours indicate age of the internal node as shown in the legend. It suggests that most of the spread happened relatively recently in the last 2000 years. Most of Africa remains white due to lack of samples in the data set in those areas.

**Figure 8:**
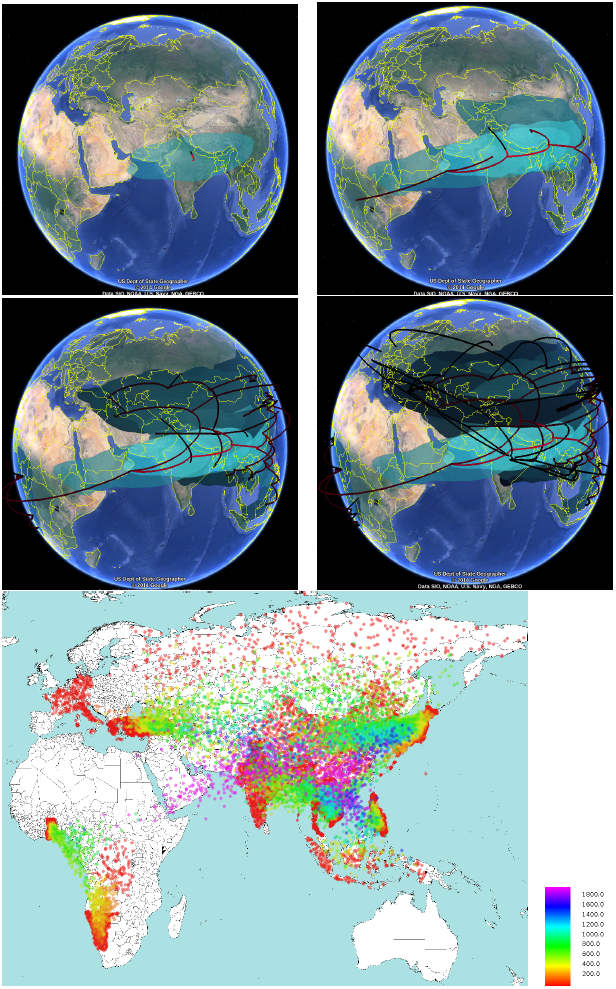
Hepatitis B in Eurasia and Africa.

## 5 Discussion

The spherical diffusion model can be used for phylogeographical analyses in situations similar to where the models of [16, 17] are used that are assuming diffusion on a plane. Furthermore, the spherical diffusion model can be used when the area of interest is large, and there is considerable distortion when the area is projected on a plane.

However, it assumes heterogeneous diffusion over a sphere. This means that, unlike the models from [16, 17], no distinction is made between possible correlations in the direction of the random walk. Furthermore, the random walk on a sphere assumes a Gaussian distribution with relatively thin tails compared to a Levy jump process on a plane [20]. Also, it assumes heterogeneous diffusion, unlike the landscape aware model [6].

The spherical diffusion model assumes locations in continuous space, which tends to be more powerful than using discrete locations. However, it also means that phylogeographical approaches based on the structured coalescent [14] cannot be applied, hence demographic developments are not captured by the geographical process.

The method is implemented in the GEO SPHERE package in BEAST 2 [5, 10], which is open source licensed under LGPL. An analysis can be set up using BEAUti, the graphical user interface for BEAST. A tutorial explaining how to use the method and set up an analysis is available from http://beast2.org/wiki/tutorials.

Results can be visualised using Google-earth after processing with SPREAD

## 6 Conclusions

We presented a new way to perform Bayesian phylogeographical analyses based on diffusion on a sphere. An approximation of the likelihood that can be calculated efficiently was presented, which can be used in an MCMC framework and is implemented in BEAST 2. The framework allows branch rate models in order to relax the strict clock assumption, as well as efficient sampling when prior information in the form of sampling regions for tip, root or monophyletic clade locations is available.

Further investigations include incorporating inheterogeneous random walks to represent more realistic diffusion processes that distinguish different rates among land and water, among forests and deserts, etc.

## Acknowledgements

Joseph Heled helped a lot with many lively discussions on this topic. This research was funded by Marsden grant (UOA1308) (http://www.royalsociety.org.nz/programmes/funds/marsden/awards/2013-awards/), a Rutherford fellowship (http://www.royalsociety.org.nz/programmes/funds/rutherford-discovery/) from the Royal Society of New Zealand awarded to Prof. Alexei Drummond. It was also funded by the Max Planck Institute.

## Appendix A: Proof of Theorem 1

*Proof.* First, we show that Equation (9) and (8) are valid for any internal node *x*_*i*_ when calculated by post-order traversal. When we reach *x*_*i*_ it can be an internal node or the root node. If *x*_*i*_ is an internal node, we distinguish between the children being leaf nodes or internal nodes. Images for Figure 1 were taken from Wikipedia.

1. *x*_*L*(*i*)_ *and x*_*R*(*i*)_ *are leaf nodes*. Then Equation (4) gives, (using *lat*_*L*(*i*)_ =*m*_*L*(*i*)_, *lat*_*R*(*i*)_ = *m*_*R*(*i*)_ for child nodes *x*_*L*_(*i*) and *x*_*R*_(*i*), and the definition of *m*_*i*_ and *ρ*_*i*_)

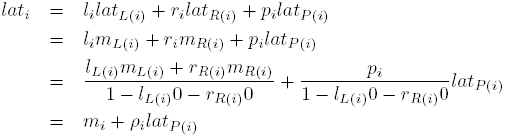
2. *x*_*L*(*i*)_ *is a leaf and x*_*R*(*i*)_ *an internal node*. Then, *lat*_*R*(*i*)_ = *m*_*R*(*i*)_+*ρ*_*R*(*i*)_*lat*_*i*_ (by post-order traversal) and *lat*_*L*(*i*)_ = *m*_*L*(*i*)_. Then Equation (4) gives,

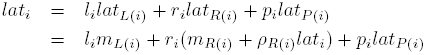

grouping *lat*_*i*_ and divide by 1 *−* 0 *− r*_*i*_*ρ*_*R*(*i*)_ gives

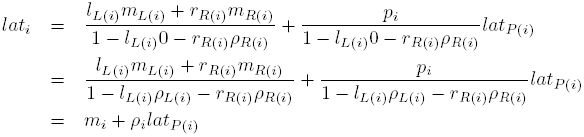
3. *x*_*L*(*i*)_ *is an internal node and x*_*R*(*i*)_ *a leaf node*. By symmetry, case 2holds.
4. *x*_*L*(*i*)_ *and x*_*R*(*i*)_ *are both internal nodes*. Then, *lat*_*L*(*i*)_ = *m*_*L*(*i*)_ + *ρ*_*L*(*i*)_*lat*_*i*_ and *lat*_*R*(*i*)_ = *m*_*R*(*i*)_+*ρ*_*R*(*i*)_*lat*_*i*_ (by post-order traversal). Now Equation (4) gives,

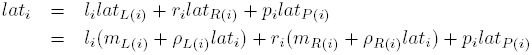

grouping *lat*_*i*_ and divide by 1 − *l*_*i*_*ρ*_*L*(*i*)_ − *r*_*i*_ρ_*R*(*i*)_ gives

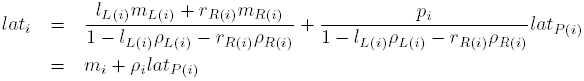

If *x*_*i*_ is the root Equation (8) follows from a similar argument, but with *ρ*_*i*_ = 0.

From equations Equations (9) and (8) together it follows that the mean can be calculated by post-order traversal to provide information for the root location (8) to be calculated, followed by a pre-order traversal to calculate locations of internal nodes using (9).

Complexity: it is easy to see the algorithm is *O*(*n*) since it takes one post-order traversal sending 2*n −* 2 messages up and one pre-order traversal calculating locations for *n −* 1 internal nodes. Each message and latitude calculation is *O*(1).

## Appendix B: HBV data

| Genbank | year | location | Genbank | year | location | Genbank | year | location |
| --- | --- | --- | --- | --- | --- | --- | --- | --- |
| AB026814 | 1998 | Japan | AB198076 | 2001 | China | AJ748098 | 2002 | Vietnam |
| AB033550 | 1988 | Japan | AB198077 | 2001 | China | AY341335 | 2003 | Greece |
| AB033554 | 1985 | Indonesia | AB198078 | 2001 | China | AY641558 | 2003 | Korea |
| AB033555 | 1984 | Indonesia | AB198079 | 2001 | China | AY641559 | 2003 | Korea |
| AB033556 | 1985 | Japan | AB198080 | 2001 | China | AY641560 | 2003 | Korea |
| AB049609 | 1996 | Japan | AB198081 | 2001 | China | AY641561 | 2003 | Korea |
| AB049610 | 1996 | Japan | AB198082 | 2001 | China | AY641562 | 2003 | Korea |
| AB106564 | 1999 | Ghana | AB198083 | 2001 | China | AY641563 | 2003 | Korea |
| AB109475 | 2001 | Japan | AB198084 | 2001 | China | AY721605 | 2004 | Turkey |
| AB109476 | 1997 | Japan | AB205010 | 1994 | Japan | AY721606 | 2004 | Turkey |
| AB109477 | 1997 | Japan | AB205118 | 2001 | Japan | AY721607 | 2004 | Turkey |
| AB109478 | 1998 | Japan | AB205119 | 2000 | Japan | AY721608 | 2004 | Turkey |
| AB109479 | 1997 | Japan | AB205120 | 2000 | Japan | AY721609 | 2004 | Turkey |
| AB110075 | 2001 | Japan | AB205122 | 2001 | Vietnam | AY721611 | 2004 | Turkey |
| AB111121 | 1998 | Japan | AB205123 | 2002 | China | AY721612 | 2004 | Turkey |
| AB111125 | 2001 | Japan | AB205124 | 2003 | Japan | AY738143 | 1997 | Germany |
| AB111946 | 1998 | Vietnam | AB205125 | 2001 | Vietnam | AY738147 | 1998 | DR Congo |
| AB112063 | 1998 | Vietnam | AB205126 | 2001 | Japan | AY796030 | 2004 | Turkey |
| AB112065 | 1998 | Vietnam | AB205127 | 2000 | Russia | AY796031 | 2004 | Turkey |
| AB112066 | 1999 | Myanmar | AB205128 | 2000 | Russia | D23680 | 1991 | Japan |
| AB112348 | 1999 | Myanmar | AB205129 | 1999 | Ghana | D23681 | 1992 | Japan |
| AB112408 | 1999 | Myanmar | AB205188 | 2000 | Ghana | D23682 | 1984 | Japan |
| AB112471 | 2002 | Thailand | AB205189 | 2000 | Ghana | D23683 | 1984 | Japan |
| AB112472 | 2002 | Thailand | AB205190 | 2000 | Ghana | D23684 | 1988 | Japan |
| AB115417 | 1996 | Japan | AB205191 | 2000 | Ghana | D28880 | 1992 | Japan |
| AB115551 | 2004 | Cambodia | AB205192 | 2000 | Ghana | DQ060824 | 1983 | Namibia |
| AB116266 | 1987 | Japan | AB212625 | 2004 | Vietnam | DQ060825 | 1983 | Namibia |
| AB117758 | 2004 | Cambodia | AB212626 | 2004 | Vietnam | DQ060826 | 1983 | Namibia |
| AB117759 | 2004 | Cambodia | AB219426 | 2002 | Philippines | DQ060827 | 1983 | Namibia |
| AB119251 | 2002 | Japan | AB219427 | 2002 | Philippines | DQ060828 | 1983 | Namibia |
| AB119252 | 2002 | Japan | AB219428 | 2002 | Philippine | DQ060829 | 1983 | Namibia |
| AB119253 | 2000 | Japan | AB219429 | 2002 | Philippines | DQ315776 | 2000 | India |
| AB119254 | 2003 | Japan | AF121243 | 1995 | Vietnam | DQ315777 | 2003 | India |
| AB119255 | 2001 | Japan | AF121244 | 1993 | Vietnam | DQ315778 | 2003 | India |
| AB119256 | 1997 | Japan | AF121245 | 1992 | Vietnam | DQ315779 | 2003 | India |
| AB120308 | 1982 | Japan | AF121246 | 1998 | Vietnam | DQ315780 | 2002 | India |
| AB126580 | 2000 | Russia | AF121247 | 1994 | Vietnam | DQ315781 | 2003 | India |
| AB126581 | 2000 | Russia | AF121248 | 1994 | Vietnam | DQ315782 | 2004 | India |
| AB179747 | 2002 | Italy | AF121249 | 1998 | Vietnam | DQ315783 | 2003 | India |
| AB188241 | 1999 | Japan | AF121250 | 1997 | Vietnam | DQ315784 | 2003 | India |
| AB188242 | 1999 | Japan | AF121251 | 1997 | Vietnam | DQ315785 | 2003 | India |
| AB188243 | 1999 | Japan | AF182802 | 1992 | China | DQ315786 | 2005 | India |
| AB188245 | 2000 | Japan | AJ344117 | 1990 | France | M57663 | 1987 | Philippines |

